# ReNette: a web-infrastructure for reproducible network analysis

**DOI:** 10.1101/008433

**Authors:** M. Filosi, S. Droghetti, E. Arbitrio, R. Visintainer, S. Riccadonna, G. Jurman, C. Furlanello

**Affiliations:** Fondazione Bruno Kessler (FBK), via Sommarive 18, I-38123 Povo (TN), Italy; Research and Innovation Centre, Fondazione Edmund Mach (FEM), I-38010 San Michele all'Adige (TN), Italy

## Abstract

**Summary:** Here we introduce a novel web-infrastructure for differential network analysis. The aim of the web-site is to provide a comprehensive collection of tools for network inference, network comparison and network reproducibility analysis. Four main processes are available through the web service: the network inference process which include 11 reconstruction algorithms, the network distance process with 3 available metrics, the network stability process which includes all the network reconstruction methods and network distances and the netwok statistic process which computes the most common measures for network characterization. We introduce here a novel infrastructure which allows the user-interface logic to be separated from computing services and the asynchronous task management. Task submission is implemented mimicking the high performance computing queue submission system which allows to run multiple jobs without affecting the front-end server.

**Availability and Implementation:** The web-site is available at https://renette.fbk.eu, the implementation is based on the django framework and Apache, with all major browsers supported. Furthermore, the whole project is Open Source under *GPLv2* and the code is available on GitHub at https://github.com/MPBA/renette for local installation.

## 1 INTRODUCTION

Given the importance network theory has gained in biology during the past decade Zanzoni *et al.* (2009); Buchanan *et al.* (2010), we provide an interface for network reproducibility called *ReNette*. The user friendly interface guide the scientist with few programming skills to run a complete network analysis pipeline on his own data. When multiple networks describing the same process under different conditions are available it is possible to compare them quantitatively with the network distance process. Furthermore it is important to study the stability of the inference process in order to understand the amount of the network variations due to random noise. This can be easily computed with *ReNette* using the network stability process. For each process we included the state-of-the art algorithms; the user can choose among them through the web-interface without installing many different packages and software. The results are presented in a synthetic form on the web page but they can also be downloaded as a zip archive and the network topology can be exported in different common formats to be used for visualization with third part software. In order to guide the user through all the available feature of *ReNette* we provide a complete tutorial for each process with an exhaustive explanation of each algorithm and parameter used. The user will follow the tutorial with a biological dataset that can be automatically loaded for each process.

## 2 FEATURES & METHODS

The *ReNette* web-infrastructure is a public service for network reproducibility analysis. We collect 11 inference methods for co-expression network, 3 quantitative network comparison metrics, 8 network statistic measures and a framework for network stability analysis.

### 2.1 Processing Algorithms

The stability framework implemented here is based on Filosi *et al.* (2014). The included inference methods are CLR, ARACNE, WGCNA, bicor and TOM Filosi *et al.* (2014), MINE and DTWMIC Albanese *et al.* (2013); Riccadonna *et al.* (2012). For WGCNA, bicor and MINE a False Discovery Rate controlled version is also implemented Filosi *et al.* (2014). The available network metrics are the Hammingdistance, the Ipsen-Mikhailovdistance and the novel HIM product metric previously proposed in Jurman *et al.* (2013). Furthermore we provide 5 network measures on the whole network and on each community detected by the community detection algorithm. A fully guided tutorial is provided on the publicly available Hepato Cellular Carcinoma gene expression dataset Budhu *et al.* (2008).

#### Implementation Details

The infrastructure is based on the *django* Model View Controller (MVC) framework with *Python* version 2.7 which interacts with the core process through the *Celery* distributed task queue. The application server was built with *apache2 web server* integrated with *uWSGI* which allows configuration on high performance computing. The core of the server is based on the *rpy2* interface which interacts with the R package *nettools*. The server structure and configuration (Fig. 1(b)) separate the user logic from the computation resources. This improves the scalability of the system, in particular computation servers can be physically separated from the front-end server. The message broker *Redis* integrated with the *Celery* queue system allows the management of asynchronous jobs, thus multiple computation servers can be allocated on demand using cloud computing.

**Fig. 1.**
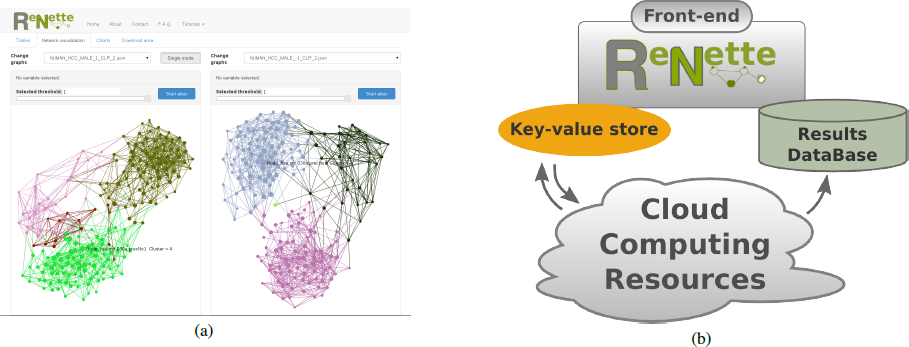
**(a)** Dual mode view for network topology comparison. The user can interact with the graph moving the sliding selector only links below the selected threshold are shown. Clicking on the single node information on node degree and node name appear. Each node is assigned to a community identified by a color.**(b)** The distributed infrastructure of the system. Computing resources are logically separated from the front-end. The database stores results and information about the jobs submitted and the communications between the front-end and the computing resources are handled by a message broker that uses a key-value store.

At the moment the system runs on a 24 cores, 72 GB of RAM and 1TB of storage machine (*georun*) with Red Hat Enterprise Linux Workstation release 6.5 with kernel 2.6.32 as operating system. The current version 1.0 needs python 2.7, django 1.6.1, rpy2 2.30, *Celery* 3.1.7, R 3.1.0, nettools 0.9.5, igraph 0.6.6 to run correctly. The server has been running for 257 days; a total amount of 436 different analysis processes for seven different users. At the moment the machine is hosted in the FBK data center on the internal trusted network and it is reachable from the web through a web-proxy which correctly redirects the allowed requests to the internal server and block the non-authorized ones. To install the system, a complete list of the dependencies together with the source code can be found on the github page https://github.com/MPBA/renette

#### Input

The network inference and network stability procedures take *CSV* files containing original measurements data such as microarray/RNA-seq datasets, all the possible relationships between columns are reconstructed by the inference algorithms. The upload of multiple files is also supported. Network distances are computed on one or more adjacency matrices as inputs (*CSV*).

## 3 RESULT VISUALIZATION

The results of the analysis are presented in different ways. For each input file the adjacency matrix is presented as a table. If the edge between two nodes is present the weight of the edge is reported in the correct position of the table. The stability process produces two more tables, one reporting the node degree variations across resampling and the other one contains the edge weight distribution. Network topologies are presented in the visualization modules. The topology for each network is visualized on the screen, node positions are assigned using the atlas algorithm as shown in Fig. 1(a). Node colors are assigned based on the communities discovered by the spinglass community detection algorithm. The inferred network can be visualized and exported in file formats supported by the IO functions in *igraph* library. For each network the degree distribution and the node weight distribution is presented in a plot. The inference and the network stability procedures return the adjacency matrix for each input file and selected reconstruction method as a *CSV* file. The network stability procedure outputs also the *netSI* Filosi *et al.* (2014) indicators in 3 different *CSV* files. The network distance returns, for each selected method, a matrix with the pairwise distances between all possible pairs of input files. The network statistics process returns a table with the measures values computed for each node. Each output can be viewed online or downloaded in a compressed folder.

## ACKNOWLEDGMENT

We thank our collaborators at the Center for Systems and Computational Biology at Wistar Institute, Philadelphia.

